# Date palm virus A: first plant virus found in date palm trees

**DOI:** 10.1101/2020.08.31.275669

**Authors:** Ayoub Maachi, Tatsuya Nagata, João Marcos Fagundes Silva

## Abstract

In this work, a novel ssRNA (+) viral genomic sequence with gene organization typical of members of the subfamily *Quinvirinae* (family *Betaflexiviridae*) was identified using high throughput sequencing data of date palm obtained from the Sequence Read Archive database. The viral genome sequence consists of 7860 nucleotides and contains five ORFs encoding for the replication protein (Rep), triple gene block proteins 1, 2, 3 (TGB 1, 2 and 3) and coat protein (CP). Phylogenetic analysis based on the Rep and the CP amino acid sequences showed the closest relationship to garlic yellow mosaic-associated virus (GYMaV). Based on the demarcation criteria of the family *Betaflexiviridae*, this new virus, provisionally named date palm virus A (DPVA), could constitute a member of a novel genus. However, considering that DPVA and GYMaV share the same genomic organization and that they cluster together on the Rep phylogenetic analysis, they could also constitute a novel genus together, highlighting the necessity of a revision of the taxonomic criteria of the family *Betaflexiviridae*.

## Text

Virus members of the family *Betaflexiviridae* have flexuous filament virions and ssRNA (+) genome, ranging 5.9-9.0 kb. The genomic RNA is presumably capped at the 5’ end and has a polyadenylated tail at the 3’ end [1]. Betaflexiviruses were reported from a diverse host range of plant species, from vegetable crops to fruit trees [1]. Date palm tree, a crop intensively cultivated and of major economic importance in Northern African and Middle East countries [2], is very robust and, until now, no virus whatsoever was reported to infect this plant species. Only African oil palm ringspot virus of this family was reported to infect palm trees (family: *Arecaceae*) [3,4], however, no reports was found about the occurrence of this virus in palm tree. Here, a plant virus sequence was found in high throughput sequencing (HTS) data of date palm trees (*Phoenix dactylifera*) from the Sequence Read Archive (SRA) public database. Thus, we report the characteristics of the genomic sequence of a novel virus belonging to the family *Betaflexiviridae*, tentatively named date palm virus A.

Raw transcriptomic data from SRA under the NCBI BioProject number PRJNA472694, used for comparison of gene expression in male and female date palm flowers, were trimmed with BBDuk v38.08 to remove adapters and low-quality sequences. The resulting reads were mapped against the date palm sequence under the RefSeq assembly accession number GCF_000413155.1 using Burrows-Wheeler Aligner (BWA) [5], then all mapped sequences were removed from the datasets. A total number of 130,973,904 filtered and trimmed paired reads with a read length of 75 bp were *de novo* assembled using MEGAHIT [6]. A total of 25,005 contigs ranging from 200 to 8523 nt long were queried against the virus RefSeq database using tBLASTx [7]. A contig of 7774 nt long was aligned to the replication protein of cherry green ring mottle virus with the amino acid (aa) sequence identity of 41.8%. Iterative read mapping function in Geneious 8.1.9 [8] allowed to extend the assembled contig to a 7860 nt long sequence, supported by 1,815,153 reads from multiple samples of date palm trees used in that study. Five ORFs were predicted using the Find ORF function in Geneious 8.1.9.

The assembled genomic sequence of the putative new virus is 7860 nt long excluding the poly(A) tail with 36.4% of GC content (accession number BK011997). The genomic organization is similar to that of viruses from the genera *Foveavirus* and *Robigovirus* of the family *Betaflexiviridae* [1]. It consists of the 5’ UTR (93 nt) followed by five ORFs encoding for structural and non-structural proteins, and a 278 nt of 3’ UTR (Fig. 1a). ORF1 is 5514 nt long (1838 aa, 212 kDa) and encodes for a polyprotein that contains the methyltransferase, helicase and replicase domains, which were all the conserved domains of *Betaflexiviridae* family. The ORF1 shares the highest nt (49.92%) and aa (40.09%) sequence identities with garlic yellow mosaic-associated virus (GYMaV). ORF2, ORF3 and ORF4 were 705 nt (235 aa; 26.6 kDa), 348 nt (116 aa; 12.7 kDa), 195 nt (65 aa; 7 kDa) long and encode the triple gene block 1, 2 and 3 (TGB1, TGB2 and TGB3), respectively, which are involved in cell-to-cell and long distance movement of the virus [9]. ORF5 encodes the coat protein (CP) gene which was 747 nt long (248 aa; 26.8kDa). The CP gene shared the highest identities (47.45% of nt and 36.40% aa sequences) with GYMaV.

**Fig. 1.**
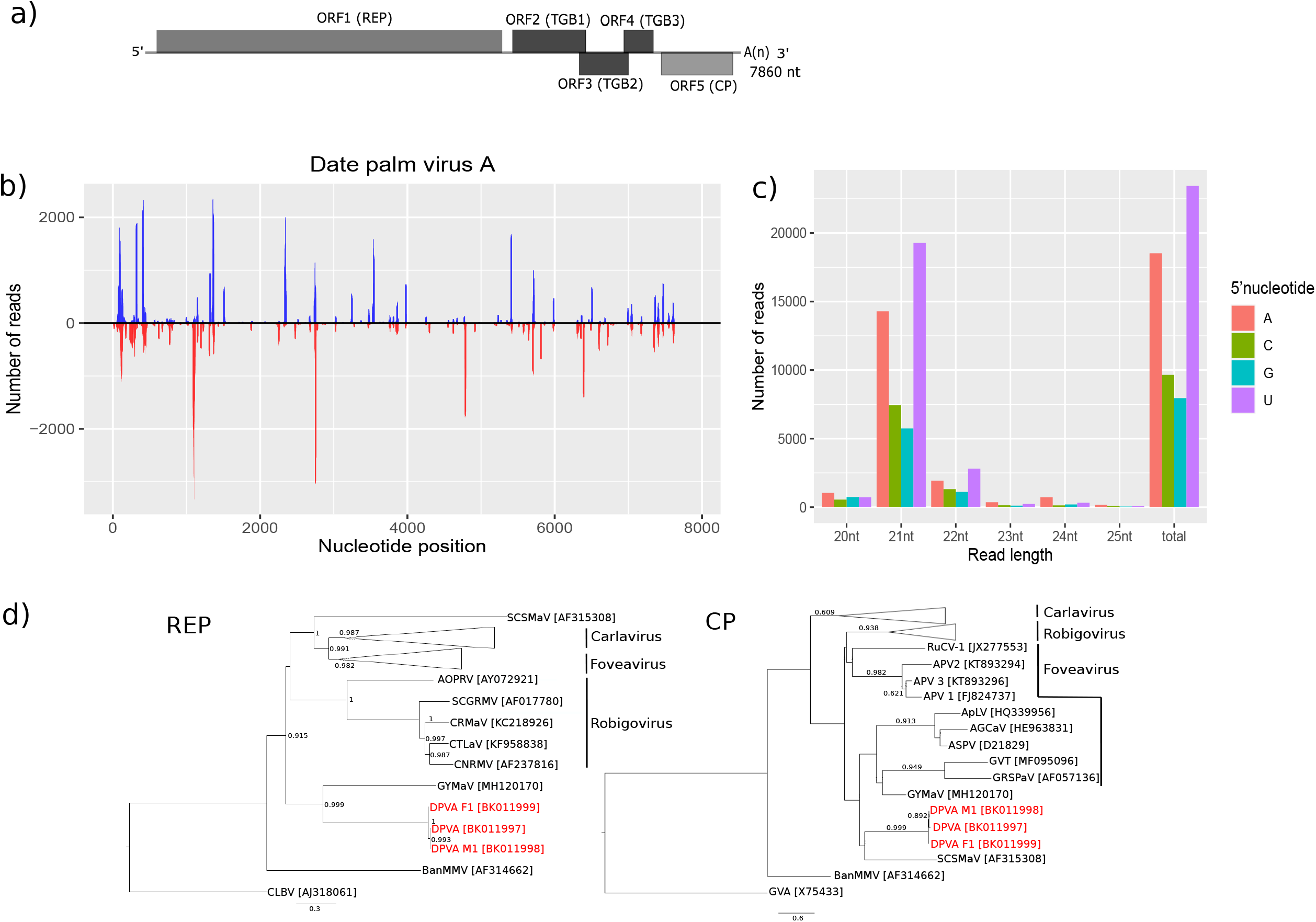
(a) Genomic organization of date palm virus A. The boxes show the positions of the five open reading frames. REP, replicase; TGB1, triple gene block 1; TGB2, triple gene block 2; TGB3, triple gene block 3; CP, coat protein; A(n), poly(A). (b) Distribution of vsRNAs along DPVA genome. (c) Number of small RNAs categories and their 5’nucleotide composition. (d) Neighbor-joining phylogenetic trees based on the amino acid sequences of the replicase protein (ReP) and the coat protein (CP) of members of the subfamily *Quinvirinae*. Trees were inferred with MEGA X. Alignments were performed with MAFFT. The newly characterized date palm A and its isolate M1 and F1 were also included. Citrus leaf blotch virus and grapevine virus A were used as outgroup for the REP and CP, respectively. SCSMaV, sugarcane striate mosaic-associated virus; AOPRV, african oil palm ringspot virus; SCGRMV, sour cherry green ring mottle virus; CRMaV, cherry rusty mottle associated virus; CTLaV, cherry twisted leaf associated virus; CNRMV, cherry necrotic rusty mottle virus; GYMaV, garlic yellow mosaic associated virus; DPVA, Date palm virus A; BanMMV, banana mild mosaic virus; RuCV-1, rubus canadensis virus 1; APV 1, asian prunus virus 1; APV 2, asian prunus virus 2; APV 3, asian prunus virus 3; ApLV, apricot latent virus; AGCaV, apple green crinkle associated virus; ASPV, apple stem pitting virus; GVT, grapevine virus T; GRSPaV, grapevine rupestris stem pitting associated virus 1.

Given that plants produce high amounts of virus-derived small RNAs (vsRNAs) upon infection, an additional dataset of ncRNA-seq from date palm trees (SRA accession numbers: SRR11553503 - SRR11553506) generated by an independent study was used to confirm whether the new putative virus indeed infects this host. Adapter sequences were determined using DNApi [10] and removed using cutadapt. Reads were mapped to the reference genome of the new putative virus sequence with BWA aln [5]. Reads containing up to one mismatch were extracted and sorted by size and 5’ terminal nucleotide with SAMtools [11]. MISIS-2 [12] was used to get the new consensus sequence from this independent data. vsRNAs were distributed along almost the complete sequence covering 63.76% of the virus sequence with an average coverage depth of 160.54 (Fig. 1b), where viral hotspots for the production of vsRNA can be observed. An enrichment for 21 and 22 nt vsRNAs was noted (78.48% and 12.02%, respectively) (Fig. 1c), suggesting that Dicer-like protein 4 plays a major role in antiviral defenses of date palm trees against this virus [13]. Loading of small RNAs in the effector complexes is determined by the identity of the 5’ end of the vsRNAs. Our analysis showed that most vsRNAs had a 5’ end uridine (Fig. 1c), indicating that DPVA-derived vsRNAs are mainly loaded into Argonaute (AGO)1, and possibly AGO10 proteins [13]. The consensus sequence obtained from that alignment showed 96% and 98.2% similarity at the nt and aa level, respectively, with the reference genome of that new virus.

The taxonomic status of the virus was interrogated vis-à-vis to the current ICTV species and genus demarcation threshold for the family *Betaflexiviridae* [1]. The nt and aa pairwise sequence identity of its Rep and CP are much below the threshold of <72 % nt and <80% aa identity for a new species. Therefore, this virus belongs to a new species in the family *Betaflexiviridae*, provisionally named Date palm virus A. The pairwise nt identity of the Rep of DPVA with GYMaV is also below the genus demarcation threshold of the family *Betaflexiviridae* (<45%), suggesting that this new virus could be a member of a novel genus in the same family. But it should be more appropriate to use aa identities for genus demarcation due to mutational saturation. In order to check the presence of DPVA in individual samples (or trees) belonging to the same BioProject, trimmed data from each sample was mapped against the previously assembled DPVA sequence. This revealed two near-complete sequences annotated as DPVA M1 (BK011998; mean coverage depth = 85.7×) and DPVA F1 (BK011999; mean coverage depth = 55.9×). Both isolates share 97.19% and 98.54% of nt identity and 96.85% and 97.90% of aa identity with the original DPVA sequence, respectively. Phylogenetic trees (Fig. 1d) were constructed with MEGA X [14] by the Neighbor-joining method with 1000 bootstrap replicates using the JTT+G substitution matrix. Phylogenetic analysis was performed with alignments of the Rep and CP with sequences available in the GenBank database that belong to the subfamily *Quinvirinae*. The DPVA isolates were positioned far from the species that are assigned to a genus of the *Quinvirinae* subfamily, and clustered closer to GYMaV in the Rep phylogeny with high support. Da Silva et al. (2019) considered that GYMaV constituted a distant evolutionary lineage in the *Betaflexiviridae* family and could be classified as a new genus in the family [15]. Both GYMaV and DPVA share similar genomic structure and evolutionary history. Therefore, these two viruses could be classified as members of a new genus, highlighting pitfalls on the current taxonomic criteria for genera in the family *Betaflexiviridae*. Inconsistencies in *Betaflexiviridae* taxonomic classification have already been noted, especially for the genus *Vitivirus* [16]. Date palm virus A is the first virus to be found in date palm trees. More studies have to be conducted to assess its epidemiology, transmission and pathogenic effect on the trees.

## Declarations

### Availability of data and materials

The new DPVA isolates described here are available at GenBank Third Party Annotation database (accessions: BK011997-BK011999).

### Author’s contributions

JMFS and TN designed and supervised the study. AM performed data analysis and wrote the manuscript. All authors contributed to the review of the manuscript.

## Compliance with ethical standards

### Research involving human participants and/or animals

This work does not contain any animal or human participants.

### Informed consent

This work does not contain any animal or human participants.

